# Adaptive discrimination of antigen risk by predictive coding in immune system

**DOI:** 10.1101/2021.12.12.472285

**Authors:** Kana Yoshido, Honda Naoki

## Abstract

The immune system discriminates between harmful and harmless antigens based on past experiences; however, the underlying mechanism is largely unknown. From the viewpoint of machine learning, the learning system predicts the observation and updates the prediction based on prediction error, a process known as ‘predictive coding’. Here, we modeled the population dynamics of T cells by adopting the concept of predictive coding; helper and regulatory T cells predict the antigen amount and excessive immune response, respectively. Their prediction error signals, possibly via cytokines, induce their differentiation to memory T cells. Through numerical simulations, we found that the immune system identifies antigen risks depending on the concentration and input rapidness of the antigen. Further, our model reproduced history-dependent discrimination, as in allergy onset and subsequent therapy. Together, this study provided a novel framework to improve our understanding of how the immune system adaptively learns the risks of diverse antigens.

## Introduction

The immune system faces the challenge of identifying unknown risks of diverse antigens and inducing proper immune responses. For harmful antigens, such as pathogens, the immune system induces strong immune responses for their elimination, whereas for harmless antigens, such as food and self-antigens, it does not lead to strong responses to prevent unnecessary inflammation. Thus, the immune system needs to discriminate between harmful and harmless antigens appropriately. Defects in this discrimination induce immune diseases, including allergies and autoimmune diseases. However, the mechanism by which the immune system distinguishes between harmful and harmless antigens upon exposure to numerous antigens remains to be understood. This study aimed to explore this field through computational modeling of T cell population dynamics using a novel concept of predictive coding.

The central organizers of adaptive immunity are T cells, each of which expresses different T cell receptors (TCRs) to specifically recognize antigens presented by antigen-presenting cells, such as dendritic cells (DCs)^1–3^. In the process of T cell differentiation, the cells responsive to self-antigens are eliminated^4–6^; however, there still remain those that are specific not only to harmful antigens but also to harmless ones. Namely, such T cells have no way of knowing whether the antigen is harmful or harmless. Nevertheless, the immune system responds strongly to harmful foreign antigens, but not to harmless ones. Therefore, we focused on the fact that the antigen specificity of T cells cannot explain the mechanism by which the immune system discriminates between harmful and harmless antigens.

Immune response is organized by the population dynamics of various cell types (Fig. 1a). It is initiated by antigen-presenting cells, such as DCs, which take in antigens and present them to T cells. Naive T (Tn) cells, with TCRs on their surface, recognize specific antigens presented by the DCs. Tn cells then differentiate into various types of T cells, such as helper T (Th) cells and regulatory T (Tr) cells, depending on cytokines, such as interleukins (ILs), in their microenvironment^7^. Th and Tr cells play distinct roles in immune responses; Th cells accelerate immune responses, leading to the elimination of antigens by activating downstream cells, such as B cells and killer T cells^7–9^, whereas Tr cells work as a brake for immune responses via the regulation of DCs and suppressive cytokines^10,11^. After the immune response, most of the induced Th and Tr cells are removed by apoptosis^12,13^; however, a small population of them differentiates into memory T cells, namely memory Th (mTh) cells and memory Tr (mTr) cells, and they persist in the body for a long time^14,15^. Upon subsequent encounter with the same antigen, memory T cells are rapidly activated, resulting in more efficient responses^16,17^. This is termed as immunological memory. Thus, the combinatorial dynamics of various types of T cells determines the intensity of immune response. In other words, the discrimination between harmful and harmless antigens must be achieved at the level of T cell population dynamics.

**Fig. 1.**
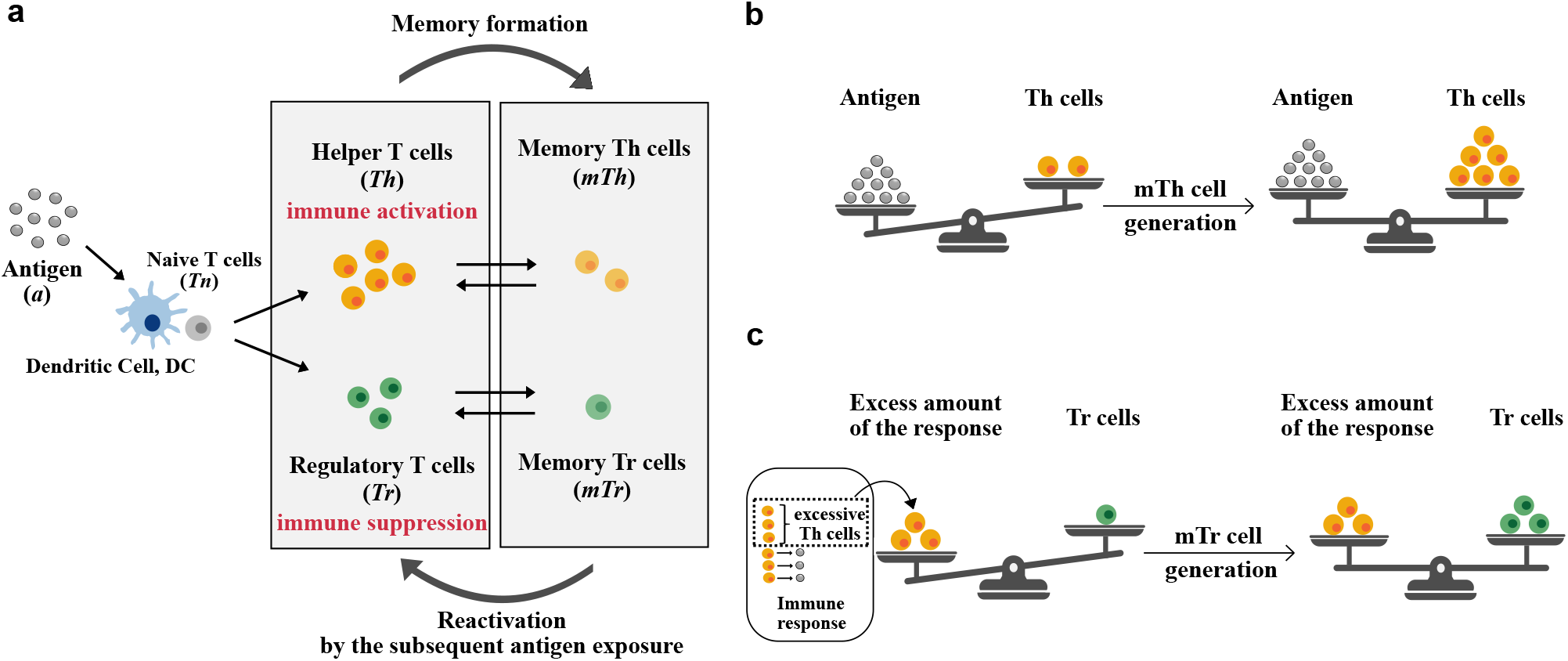
Scheme of the predictive immune memory model. **(a)** Population dynamics model of immune T cells in response to antigen input. The model includes the differentiation of Tn cells into Th and Tr cells by antigen-presenting cells such as dendritic cells (DCs), the differentiation of Th and Tr cells into memory T cells, and the reactivation of memory T cells into T cells upon subsequent exposure to antigens. **(b, c)** Predictive coding-based immunological memory formation. (b) Generation of mTh cells. mTh cells are generated based on the prediction error *e_h_*|*a* – *m_h_Th*|_+_. In other words, the production of mTh cells is induced when the concentration of antigens is excessive compared to that of Th cells in order to efficiently eliminate antigens. (c) Generation of mTr cells. mTr cells are generated based on the prediction error *e_r_*|*g*(*Th*) – *a* – *m_r_Tr*|_+_. In other words, the production of mTr cells is induced when the excess amount of response, evaluated by the difference between the intensity of Th cell activation (*g*(*Th*)) and antigen concentration, is larger than the concentration of Tr cells in order to prevent unnecessary inflammation.

Identification of the risk of antigens is not always constant and varies in a context-dependent manner. The most representative example is the onset and therapy of allergy, which is defined as an excessive response to harmless antigens, including pollen and mites. Although allergens, defined as the substances which cause allergy, are initially regarded as harmless in our body, response to them can intensify upon repeated exposures, leading to allergic symptoms. Such a change in responsiveness indicates that immune discrimination of allergens can change from harmless to harmful. Furthermore, allergic symptoms, immune responses to allergens, can be weakened by allergen immunotherapy^18–20^, in which a small amount of allergen extract (not enough to cause symptoms) is repeatedly administered to the patients; after the therapy, allergic symptoms do not occur even when patients are exposed to a large amount of the allergen. This means that discrimination can be reversed from harmful to harmless through immunotherapy. Thus, the immune system adaptively changes discrimination depending on the temporal history of antigens. Experimentally and clinically, immunotherapy has been reported to induce regulatory cell populations, such as Tr cells, and suppressive cytokines, such as IL-10^21–23^. However, the mechanisms by which immune discrimination is adaptively updated by antigen experience largely remains unclear.

The immune system can be viewed as an adaptive learning system that updates the discrimination of antigen risk. To induce the most appropriate responses, the immune system needs to predict and prepare for the subsequent invasion of antigens by the formation of memory cells. From the perspective of machine learning theory, a more accurate prediction is achieved by repeated observation and prediction, in which the prediction is updated based on prediction error, which is the difference between observation and prediction. This concept, called ‘predictive coding’, was originally proposed in neuroscience^24^, and has been widely accepted as a guiding principle for understanding learning systems, such as brain and artificial intelligence^25–27^. In this study, we adopted this concept to understand the immune system as a learning system. We hypothesized that Th and Tr cells predict the risk of antigens and excessive response, respectively, and their predictions can be updated by prediction errors via the production of memory T cells.

Based on the idea of predictive coding, this study aimed to reveal how the immune system discriminates between harmful and harmless antigens and how it changes response depending on the history of antigens. We developed a mathematical model of antigen-induced T cell population dynamics named “the predictive immune memory model”. By simulating the model, we demonstrated that the immune system can discriminate between harmful and harmless antigens using the predictive coding mechanism in an antigen concentration- and input rapidness-dependent manner. The model also demonstrated antigen history-dependent immune discrimination, as seen in the onset and therapy of allergy. Furthermore, we found that the dose-response of T cell activation does not affect the immunotherapy outcome but changes its persistence upon an additional higher exposure to allergens.

## Results

### Model for T cell population dynamics

To examine how the immune system discriminates between harmful and harmless antigens at the level of T cell population, we developed a mathematical model for the population dynamics of T cells, and named it “the predictive immune memory model” (Fig. 1a). The model consists of Th, Tr, and their memory cells. Th and Tr cells are generated by the differentiation of Tn cells and activation of memory T cells, as shown below.

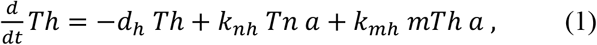

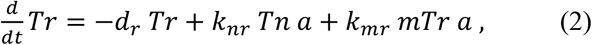

where *Th* and *Tr* represent the populations of Th and Tr cells, respectively; *mTh* and *mTr* represent the populations of memory Th and memory Tr cells, respectively; *Tn* indicates a positive constant which represents the population of Tn cells; *a* represents the concentration of antigen input; *d_i_*, *k_ni_*, and *k_mi_* (*i*. ∈ {*h*, *r*}) indicate the rates of death due to apoptosis, differentiation from Tn cells, and production of T cells from memory T cells, respectively. Memory T cells differentiate from Th and Tr cells as

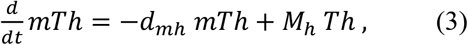

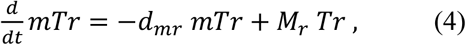

where *d_mh_* and *d_mr_* indicate the death rates of mTh and mTr cells, respectively. We regarded their death rates as zero in the time span of our simulations due to longevity of memory T cells (*d_mh_* = *d_mr_* = 0). Note that *M_h_* and *M_r_* are not constant parameters but are situation-dependent, following the idea of predictive coding (see the next section for details). In this model, we defined the intensity of response *R*, which is positively and negatively regulated by Th and Tr cells, respectively, as

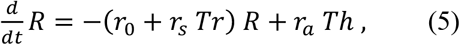

where *r*_0_ indicates a positive constant, which causes the convergence of *R* to zero in absence of Th and Tr cells; *r_a_*, and *r_s_* indicate the activation rates by Th cells and suppression rates by Tr cells, respectively.

### Predictive coding scheme

We have introduced the concept of predictive coding under the hypothesis that the immune system predicts the next antigen invasion and its consequent inflammation in an antigen experience-dependent manner. More specifically, the predictive coding scheme states that Th and Tr cells are predictors of the antigen amount and excess amount of immune response, respectively, and that their predictions are updated based on prediction errors via the formation of mTh and mTr cells.

Since Th cells are the control center to achieve antigen elimination by inducing downstream reactions, they must be adequately controlled depending on the change in antigen concentration; when the concentration of antigens is excessive compared to that of Th cells, more Th cells need to be generated in order to completely eliminate the antigens (Fig. 1b). Accordingly, the production rate of mTh cells can be described by

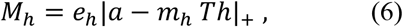

where *e_h_* and *m_h_* indicate positive constants, and |*x*|_+_ represents ramp function (i.e., |*x*|_+_ = 0 (*x* < 0), *x* (*x* ≧ 0)). Note that *M_h_* is the prediction error of antigen concentration, since *a* and *m_h_Th* represent the observation and prediction of the antigen concentration, respectively. Thus, mTh cells are upregulated by the prediction error *m_h_* (Eq. 6).

On the other hand, Tr cells play an important role in the prevention of excessive immune responses. Thus, their amount should be regulated based on the intensity of response; when the excess amount of immune response is larger than the concentration of Tr cells, more Tr cells need to be generated in order to suppress the excessive immune responses (Fig. 1c). Therefore, the production rate of mTr cells can be described by

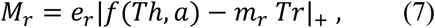

where *e_r_* and *m_r_* indicate positive constants, and *f*(*Th*, *a*) = *g*(*Th*) – *a* represents the excess amount of immune response compared to the antigen concentration. Here, we assumed that Tr cells evaluated the level of Th cell activation by *g*(*Th*) = *A_max_Th*/(*Th* + *K*), where *A_max_* and *K* indicate positive constants. Note that *M_r_* is the prediction error of the excess amount of immune response, since *f*(*Th*, *a*) and *m_r_Tr* represent the observation and prediction of the excess amount of immune response, respectively. Thus, mTr cells were upregulated by the prediction error *M_r_* (Eq. 7).

As an implementation of memory formation based on predictive coding, we assumed that the calculation of predictive coding can be achieved by cytokines. Cytokines secreted from immune cells determine their differentiation and proliferation under communication across various types of immune cells^7,28^. Their function has been vigorously explored; however, their quantitative function, where the ensemble of their concentration determines the differentiation and proliferation of immune cells, has not been elucidated. In this study, we regarded cytokines as the medium for transmitting such quantitative information. Specifically, the amounts of Th and Tr cells were coded by the concentration of cytokines secreted by themselves, whereas the amounts of antigens were coded by cytokines secreted from antigen-presenting cells, such as DCs and macrophages. Based on such information-carrying cytokines, we hypothesized that the information from each kind of cytokines is integrated into T cells, and prediction errors are computed by intracellular signal transduction in T cells. The parameters used in the numerical simulations are provided in Table S1.

### Concentration-dependent discrimination between harmful and harmless antigens

To examine the difference between harmful and harmless antigens for the immune system, we focused on the effect of antigen concentration on immune response. We simulated the model with high and low concentrations of antigen input (Fig. 2a and b), and found that the steady exposure of high and low concentrations of antigens caused more accumulation of mTh and mTr cells, respectively. At high antigen concentrations (Fig. 2a), mTh cells were generated until the prediction error *e_h_*|*a* – *m_h_Th*|_+_ was minimized to zero. mTr cells were not generated, since the prediction error *e_r_*|*f*(*Th*, *a*) – *m_r_Tr*|_+_ was always zero (left panel in Fig. 2c). Therefore, the intensity of immune response *R* converged to a high level. On the other hand, at low antigen concentrations (Fig. 2b), mTh cells were produced, similar to the exposure of high antigen concentration. mTr cells were generated more, since the prediction error *e_r_*|*f*(*Th*, *a*) – *m_r_Tr*|_+_ was positive and the generation of mTr cells continued until the prediction error was minimized to zero (right panel in Fig. 2c). Therefore, the intensity of immune response was low. To summarize the immune responses depending on antigen concentrations, there was the threshold of the antigen concentration 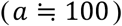, indicating that immune responses were specifically suppressed under low concentration of antigen exposures (Fig. 2d). These results suggested that the immune system with predictive coding discriminates between harmful and harmless antigens based on antigen concentration.

**Fig. 2.**
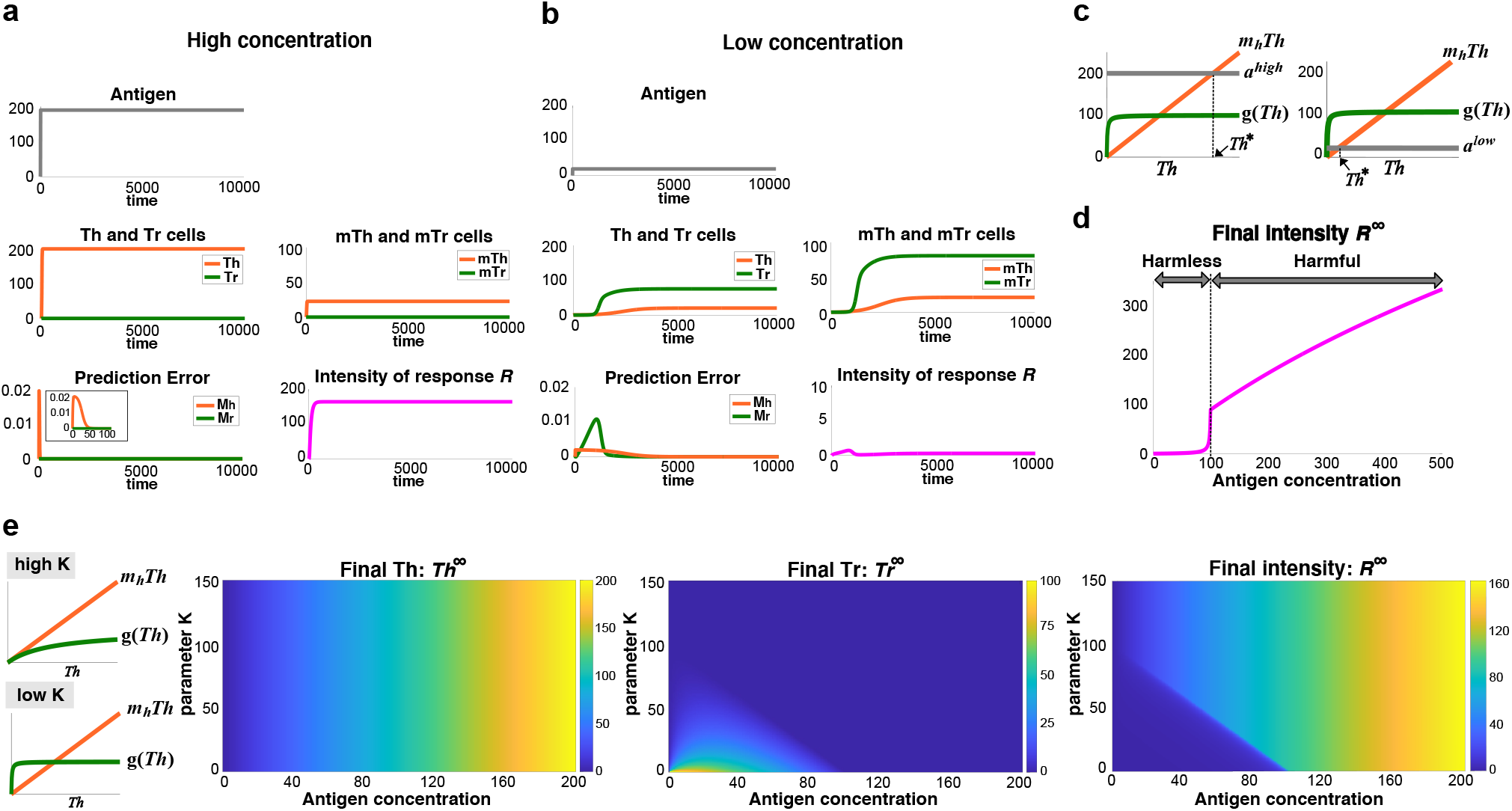
Antigen concentration-dependent immune discrimination. **(a, b)** Immune responses simulated with the exposure of (a) high and (b) low concentration of antigens. **(c)** Diagram for predictive coding-based memory formation. Antigen concentration is predicted by Th cells (orange line). When antigen concentration is observed (gray line), positive prediction error (*a* – *m_h_Th*) increases mTh cells, until *Th* converges to *Th*\* at the intersection of orange and gray lines. On the other hand, Tr cells are assumed to predict the excess amount of immune response. The green line indicates the intensity of Th cell activation evaluated by Tr cells. The excess amount of immune response is evaluated by the difference between this intensity of Th cell activation and observed antigen concentration. When antigen concentration is observed (gray line), positive prediction error between observed excess amount of immune response (*g*(*Th*) – *a*) and its prediction (*m_r_Tr*) increases mTr cells, until the prediction error converges to zero. The left and right panels show the diagram for the case where observed antigen concentration is high and low, respectively. **(d)** Change in intensity of immune responses depending on antigen concentrations. Convergence value of the intensity *R* is plotted upon the steady exposure to each antigen concentration. **(e)** Steady state responses of Th and Tr populations and the immune intensity *R* depending on antigen concentration and a parameter *K* in *g*(*Th*). Left panels visually represent how the parameter *K* in *g*(*Th*) changes the diagram for predictive coding-based memory formation as in (c), where we can regard *m_h_Th* (orange line) and *g*(*Th*) (green line) as the evaluation of Th cell activation in mTh and mTr cell generation, respectively.

Next, we examined the conditions necessary for the immune system to properly distinguish between harmful and harmless antigens depending on their concentration. We performed antigen concentrationdependent simulations by varying the parameter *K*, which regulated the Tr cell-estimated level of Th cell activation in mTr generation (left panels in Fig. 2e). We found that antigen discrimination could be achieved only with low *K*, in which immune responses were specifically suppressed under low concentration of antigen exposures (Fig. 2e). Because Tr cells underestimated and overestimated the level of Th cell activation in low antigen concentration with high and low *K*, respectively (left panels in Fig. 2e), this result suggested that suppressive immune responses at low concentration of antigens can be achieved by the overestimation of Th cell activation in mTr generation compared to the estimation in mTh generation at low antigen concentration.

### Input rapidness-dependent discrimination between harmful and harmless antigens

We focused on the rapidness of antigen input as another possible factor for discrimination between harmful and harmless antigens. We simulated the model in response to antigen inputs with different time constants (Fig. 3a and b). Similar to that seen in Fig. 2a, the intensity of immune response was high upon rapid exposure to high concentrations of antigens (Fig. 3a). However, when the concentration of antigens increased slowly, eventually reaching a high concentration, the intensity of immune response became weaker (Fig. 3b). This was because the slowly increasing antigen input enabled the immune system to have a longer experience of low concentration of antigens before reaching a high concentration, which caused positive prediction error in mTr generation followed by the production of mTr cells.

**Fig. 3.**
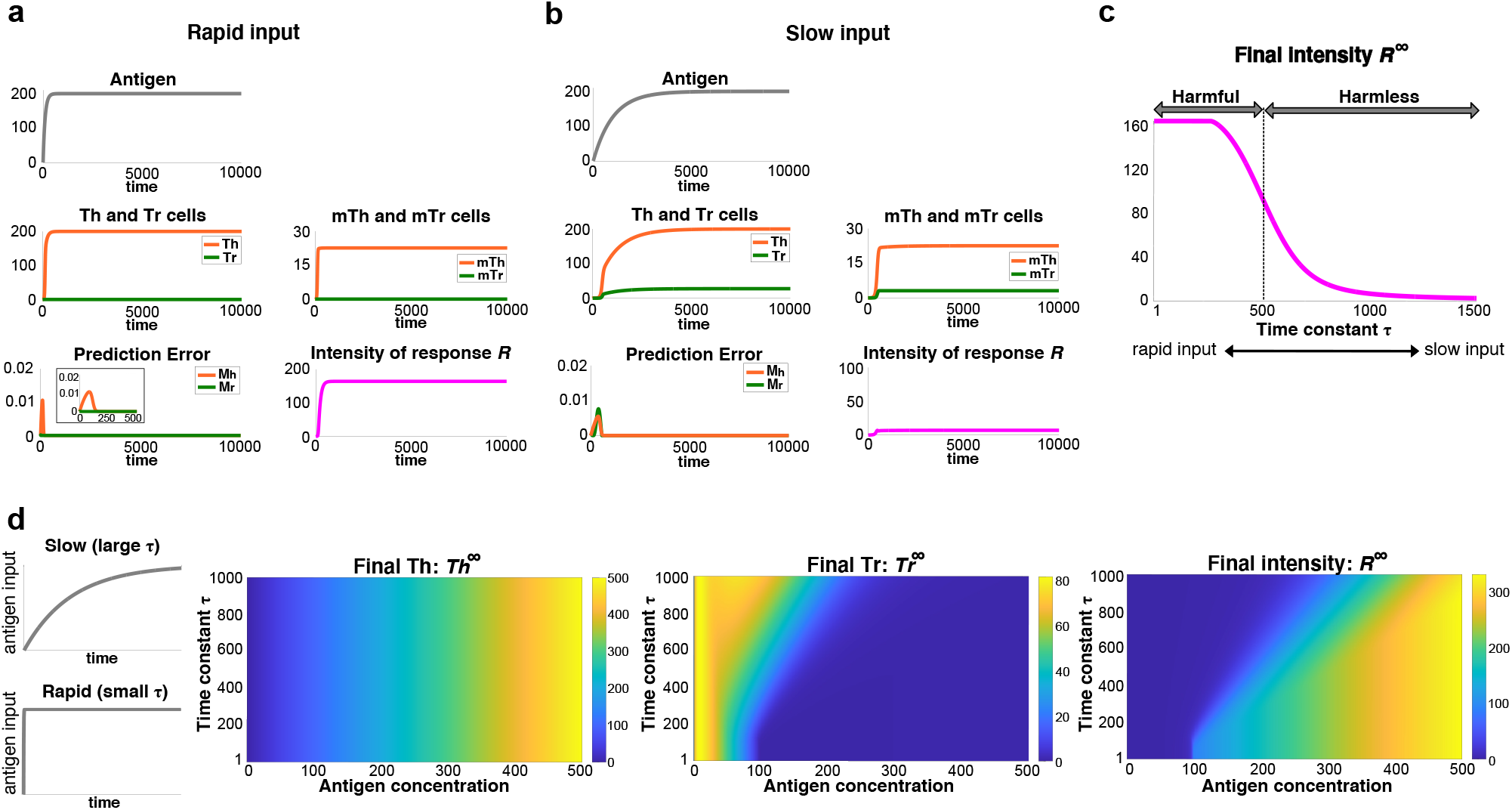
Antigen input rapidness-dependent immune discrimination. **(a, b)** Immune responses to (a) rapid and (b) slow inputs of high antigen concentration. Antigens were administered as *a*(*t*) = *a*_0_(1 – *e*^-*t*/*τ*^), where *τ* indicates the time constant. **(c)** Change in intensity of immune responses depending on time constants of antigen administration. Convergence value of the intensity *R* is plotted at each time constant. High antigen concentrations (*a*_0_ = 200) were administered. **(d)** Steady state response of Th and Tr populations and the immune intensity *R* depending on antigen concentration *a*_0_ and time constant *τ*. Left panels visually represent how time constant *τ* affects rapidness of antigen inputs.

Next, we examined input rapidness-dependent immune responses under exposure to the same high concentration of antigens with different input time constants, and found that there was a threshold of time constant for discrimination between harmful and harmless antigens 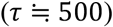 (Fig. 3c). This result showed that even when the final concentration was high, the immune system could recognize the antigens with slow input as harmless. To summarize the results, we examined the immune responses depending on both antigen concentration and its input rapidness (Fig. 3d). We found that low concentrations of antigens induced suppressive responses independent of their input rapidness. On the other hand, high concentrations of antigens induced responses with different intensities depending on their input rapidness; the immune system caused strong responses to rapidly increasing antigens while it caused suppressive responses to slowly increasing antigens. Together, these results suggested that the immune system discriminates between harmful and harmless antigens based on their input rapidness as well as their concentration.

### History-dependent discrimination between harmful and harmless antigens

Discrimination between harmful and harmless antigens can change throughout our life span. For example, at the onset of allergy, discrimination of the same antigen changes from harmless to harmful, whereas its discrimination can be reversed by allergen immunotherapy. To examine the mechanism of antigen history-dependent changes in immune discrimination, we simulated the immune responses to successive but different patterns of antigen exposure (Fig. 4a). Specifically, we applied rapid exposure of high concentrations of antigens inducing allergy, followed by exposure to low concentrations of antigens, as allergen immunotherapy, and subsequently, rapid exposure to high concentrations of antigens again. The final input was provided to examine the effect of immunotherapy.

**Fig. 4.**
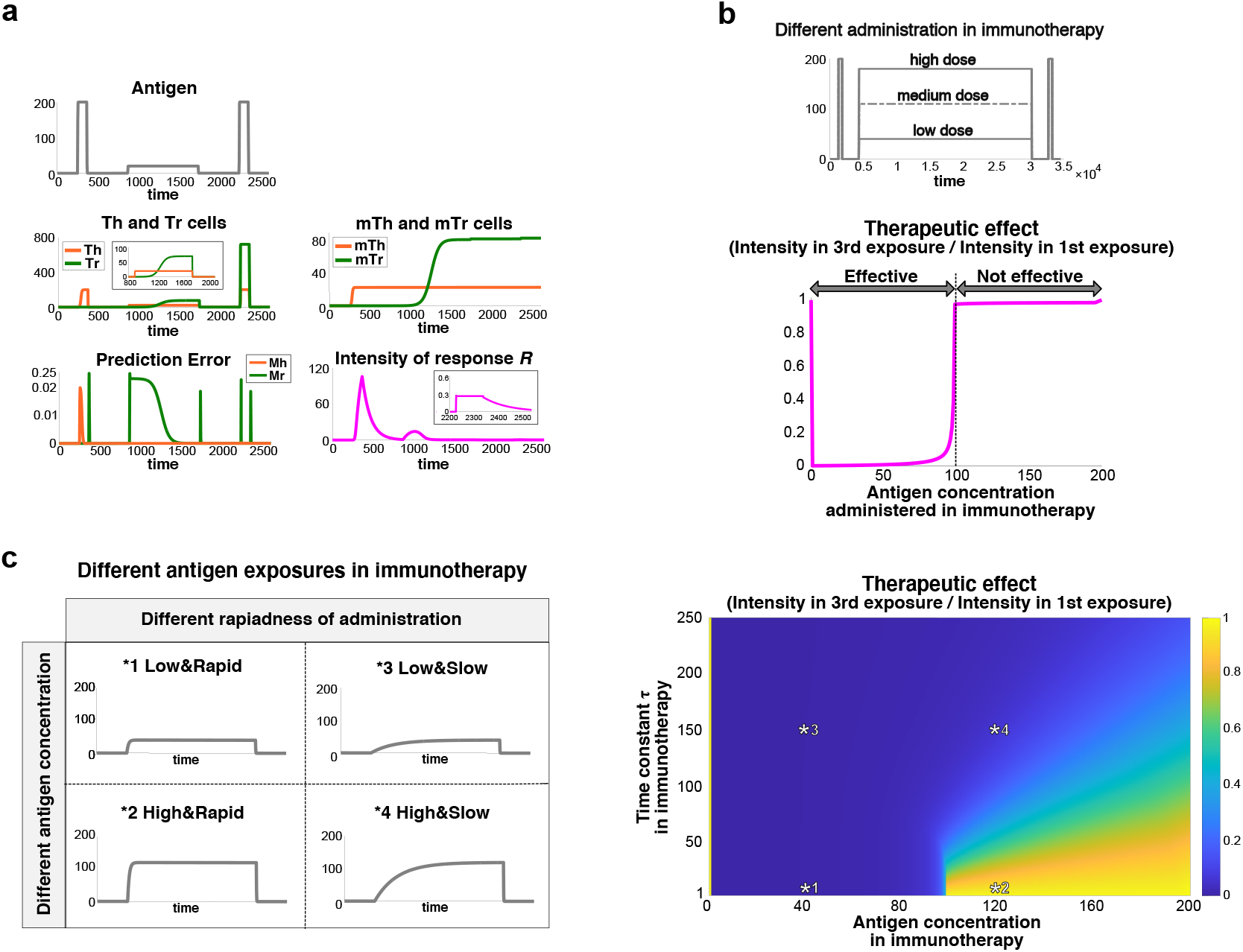
Antigen history-dependent immune discrimination in immunotherapy. **(a)** Temporal change in immune responses to a series of different antigen inputs. The first antigen input was high enough for the induction of allergy, the second one was applied for immunotherapy, and the third one was for checking the therapeutic effect. Insets show enlarged view of the populations of Th and Tr cells during immunotherapy and the intensity of response during the third antigen input. **(b)** Therapeutic effects depending on antigen concentration administered during immunotherapy. Therapeutic effect was evaluated by the ratio of maximum immune intensity *R* in the third antigen input to that in the first antigen input. A ratio smaller than one means success of the therapy. Upper panel shows the schedule of antigen inputs with different doses in immunotherapy. **(c)** Effect of immunotherapy depending on antigen concentration and rapidness of antigen inputs during immunotherapy. The left diagram shows examples of antigen input in immunotherapy with different concentrations and rapidness, corresponding to asterisks in the right panel.

After the first exposure to high concentrations of antigens, a strong immune response was induced due to the positive prediction error in mTh cell generation, as shown in antigen concentration-dependent and input rapidness-dependent discrimination. Upon exposure to low concentration of antigens thereafter, more Th cells were produced than Tr cells at the initiation of therapy, due to the accumulated mTh cells. In contrast, Tr cells were gradually generated since low concentration of antigens achieved a positive prediction error in mTr cell generation. Accordingly, exposure to low concentrations of antigens for a certain period of time enabled the accumulation of mTr cells. Therefore, even when the immune system was exposed to high concentrations of antigens again, more Tr cells were generated and the intensity of immune response became weak. This result means that the simulation successfully reproduced the immune response at the onset of allergy and the effect of allergen immunotherapy. In summary, the immune system could discriminate between harmful and harmless antigens upon the first exposure of antigens in an antigen concentration- and input rapidness-dependent manner, and its discrimination could adaptively change due to memory formation, based on predictive coding, in an antigen history-dependent manner.

Furthermore, we examined how therapeutic strategies affect the effect of immunotherapy by evaluating it by the ratio of maximum intensity *R* in response to the antigen input after the therapy to that before the therapy. We found that immunotherapy was effective only when a low antigen dose was administered for the therapy (Fig. 4b), which is consistent with the accumulation of mTr cells at low antigen concentrations as shown in Fig. 2. In addition, we examined the effect of both antigen concentration and input rapidness in immunotherapy on the therapeutic effect (Fig. 4c), and found that low concentration and/or slow input enabled effective immunotherapy. These findings could explain the validity of therapeutic strategies currently used in many clinical treatments, in which the antigen administration is initiated at low dose, and then gradually increased in the early phase of immunotherapy, mainly to prevent the patients from generating allergic symptoms during the therapy^29–31^.

### The property of T cell activation affected the history-dependent discrimination

The model has so far assumed that T cells are linearly activated by antigen concentration. However, antigen concentration-dependent activation of T cells might follow various types of activation patterns, since the difference in ligands and its consequent difference in binding properties to TCRs largely affect the T cell activation potency^32–34^. Thus, we examined the effect of dose-response types of T cell activation on immune discrimination (see Methods). Here, we simulated the model with three types of dose-response curves, namely linear, sigmoidal, and step-like types, for both Th and Tr cells (Fig. 5a, d, and g). We found that different doseresponse types of T cell activation induced different accumulation of mTh cells depending on antigen concentrations (top panels in Fig. 5b, e, and h). mTh cells accumulated to an approximately constant value, independent of the antigen concentration, in case of linear dose response, while it transiently peaked and then constantly increased with the antigen concentration in cases of the sigmoidal and step-like dose response. In contrast, the three dose-response types of T cell activation did not show a critical difference in the accumulation of mTr cells (middle panels in Fig. 5b, e, and h), and antigen concentration-dependent discrimination was achieved in all dose-response types (bottom panels in Fig. 5b, e, and h). Thus, the types of dose-response, or the properties of T cell activation, largely affected the accumulation of mTh cells.

**Fig. 5.**
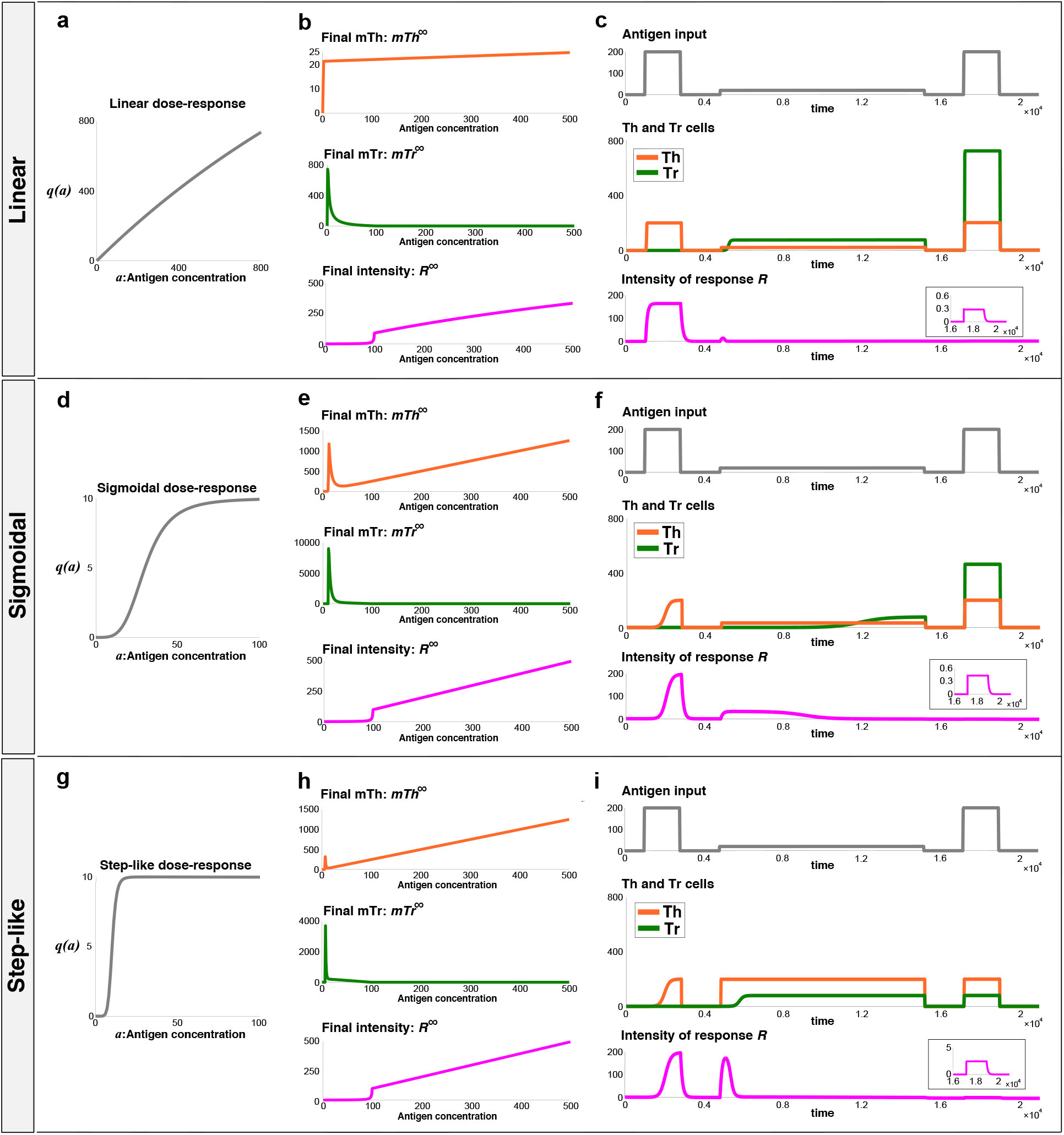
Robustness of allergen immunotherapy effect against the property of T cell activation. **(a, d, g)** Three types of dose-responses of T cell activation to antigen concentrations. *q*(*a*) represents the dose-response curves of Th and Tr cell activation. **(b, e, h)** Change in mTh and mTr cell accumulation and the intensity of immune responses, depending on antigen concentrations. The convergence values of mTh and mTr cell populations and the intensity are plotted upon steady exposure to each antigen concentration. **(c, f, i)** Temporal change in immune responses before, during, and after the immunotherapy (as in Fig. 4). Insets show enlarged view of the intensity of responses during the third antigen inputs.

Next, we examined the effect of dose-response types of T cell activation on history-dependent discrimination by simulating immunotherapy, as shown in Fig. 4 (Fig. 5c, f, and i). We found that immunotherapy was successfully achieved in all three dose-responses, since the administration of low concentrations of antigens caused a positive prediction error in mTr cell generation (details in Fig. S1). However, in the step-like dose-response, the immune intensity was high at the initiation of immunotherapy due to the production of Th cells from mTh cells, which did not depend on the antigen concentration above the threshold. This implied that some patients with allergy, with a step-like dose-response, might exhibit allergic symptoms at the early stage of therapy.

From clinical viewpoint, it is important whether the effect of therapy is persistently maintained against subsequent exposures to antigens. To examine persistence of the therapeutic effect, we considered the case where, after the therapy, patients were exposed to an additional higher concentration of antigens and subsequent lower concentration of antigens (top panels in Fig. S1). The final antigen input was applied to quantify the persistence of the therapeutic effect. We found that immunotherapy was effective in all combinations of dose-response types for Th and Tr cells (Fig. S1 and Fig. S2a). However, its effect was persistently maintained against additional exposures to higher concentrations of antigens only when the doseresponse type of Th cells was linear (Fig. S1 and Fig. S2b). These results indicated that long-term effect of immunotherapy can be determined by the dose-response type of Th cell activation. This may explain the heterogeneous effects of immunotherapy across patients, as seen in cases where some patients have relapsed allergic symptoms after discontinuing immunotherapy^29–30^.

## Discussion

In order to understand how the immune system discriminates between harmful and harmless antigens despite their diversity, we developed a generalized model that does not assume any prior information on whether each antigen is harmful or harmless. We introduced predictive coding into T cell population dynamics, opening up a new concept of immunology. Specifically, we developed a mathematical model of T cell population dynamics under the hypothesis that Th and Tr cells are predictors of the risk of antigens and excessive immune response, respectively, and their responses are regulated by prediction errors via memory T cell generation. This predictive immune memory model led to both antigen concentration- and input rapidness-dependent discrimination between harmful and harmless antigens. In addition, our model showed that such a discrimination can change in an antigen history-dependent manner, as seen in the onset and therapy of allergy. To the best of our knowledge, this is the first learning system-based model of discrimination between harmful and harmless antigens by the immune system facing diverse antigens.

Phenomenologically, harmful antigens usually originate from bacteria and viruses, and show a rapid exponential increase in their population once they invade the body. In contrast, harmless antigens, such as food, do not sharply increase in amount inside the body, but are expected to change gradually over time. Such distinct characteristics of harmful and harmless antigens can be distinguished by antigen concentration- and input rapidness-dependent discrimination (Fig. 2 and 3). Clinically, immune discrimination for the same antigen is known to change over time; for example, the onset of allergy due to exposure to high concentrations of antigens and its remission through allergen immunotherapy. This can be represented by antigen history-dependent discrimination (Fig. 4).

In this study, we introduced various types of the dose-response of T cell activation into the model, since the difference in ligands and its consequent difference in binding properties to TCRs largely affect the T cell activation potency^32–34^. Our results showed that the types of dose-response of T cell activation influenced antigen history-dependent changes in immune response, as seen in allergen immunotherapy and the subsequent recurrence (Fig. 5, S1, and S2). Overall, these results suggested that various dose-responses of T cell activation cause heterogeneity in the immune responses of individuals and/or types of antigens. In fact, some allergic patients acquire persistent remission of the symptoms by immunotherapy while others exhibit the symptoms again despite the therapy^29,30^. Moreover, allergens, such as food and bee venom, sometimes induce lethal symptoms while others, such as pollens, rarely do so^35^.

### Biological relevance of the model

Our model is a minimal model that describes essential immune processes at the level of T cells, including antigen presentation by DCs, differentiation from Tn cells to T cells, reactivation of memory T cells to T cells and memory formation. Although there are several subtypes of Th cells, such as Th1, Th2, and Th17, which induce different downstream responses, we integrated these subtypes into a single Th cell population, since all Th cell subtypes have almost the same role in terms of elimination of target antigens via different mechanisms.

Downstream of Th and Tr cells, various types of cells are involved, such as killer T cells, B cells, macrophages, neutrophils, eosinophils, basophils, natural killer T cells, and mast cells. Although we need to consider these various immune cells to discuss the whole immune response, in principle, each subtype of Th cells facilitates the activation of these downstream cells while Tr cells suppress the same^7,9^. Therefore, our model simply assumed that the intensity of immune responses can be evaluated only by the effects of Th and Tr cells.

Some immune cell populations that eliminate antigens, such as killer T cells, Th cells, B cells, and natural killer T cells, are known to persist in the body for a long time to prepare for the second infection after the first antigen experience by natural infection and vaccination^36^. It has also extensively been focused on whether regulatory immune cell subsets, such as Tr cells, generate memory populations after antigen exposures^37^. Due to the lack of memory-specific phenotypic markers for the identification of these populations, it remains controversial whether distinct memory subsets contribute to the persistence of immunosuppressive effects^38,39^. However, several studies have defined memory Tr cells and revealed their characteristics as memory populations^13,40^. Hence, our model included memory Tr population as one of the possible implementations of regulatory memory formation.

Cytokines play an important role in the induction of immune responses. Many experimental studies have identified a variety of cytokines and their functions related to the differentiation and proliferation of T cells^7,28^. For example, IL-12 and interferon *γ* (IFN*γ*) induce the differentiation of Th1 cells, and IL-2 and IFN*γ* secreted from them activate killer T cells and mononuclear phagocytes including macrophages, respectively. On the other hand, IL-4 and IL-2 induce the differentiation of Th2 cells, and IL-4 secreted from them is involved in allergic inflammation. In addition, transforming growth factor-*β* (TGF-*β*) is a critical cytokine for the differentiation of induced Tr cells, and IL-10 secreted from these cells exerts an immunosuppressive effect. Although the molecular basis of cytokines has been extensively investigated, their information basis, i.e., the types of information they encode, have not been explored much. In this study, we assumed the production of memory T cells based on predictive coding. For implementation, we regarded cytokines as the media for transmitting quantitative information. Specifically, the amounts of Th and Tr cells could be coded by the concentration of cytokines secreted by themselves, whereas the amounts of antigens could be coded by the concentration of cytokines secreted from antigen-presenting cells, such as DCs and macrophages. Based on such information-carrying cytokines, we hypothesized that prediction errors in predictive coding can be computed by intracellular signal transduction in T cells. This hypothesis can be validated by quantifying the time series of T cell populations with each TCR and cytokines in future experiments.

### Comparison with previous models

Several studies have reported computational models of immune dynamics. Different models of T cell population dynamics have focused on allergen immunotherapy. In one model, allergen immunotherapy was represented by prolonged activation of Tr cells with a large time constant^41^. In another model, the effect of allergen immunotherapy was represented by a transition from a Th2 cell-dominant state to a Tr cell-dominant state^42^. However, effect of the therapy spontaneously disappeared after elimination of antigens due to the absence of explicit T cell memory.

Immune discrimination had earlier been assessed by various mathematical models. Sontag had modeled the interaction between T cell population and antigens, such as pathogens and tumor cells^43^; this study revealed immune discrimination based on dynamic features of antigen presentation, such as the growth rate of antigens. In addition, Pradeu et al. had proposed the discontinuity theory stating that discontinuous (sudden or intermittent) exposures to antigens induce vigorous immune responses, whereas progressive and persistent exposures induce weak responses^44^. These studies were consistent with our results in terms of immune discrimination being independent of antigen type; however, they lacked immunological memory formation.

### Future perspectives

In this study, our model revealed immune discrimination based on universal information about all kinds of antigens, such as their concentration and input rapidness, and reported temporal changes based on the history of antigen exposures. However, our current model considered antigen-induced responses of T cells specific to only one kind of antigens and it did not include antigens which undergo self-renewal and can be eliminated by immune system, such as pathogens. In previous studies, Tanaka and colleagues have focused on the onset and therapy of atopic dermatitis and developed a mathematical model describing the interaction of pathogens, skin barrier integrity, and innate/adaptive immune system. They revealed different phenotypes in patients, derived from certain parameters (genetic risks), and suggested an effective treatment strategy based on optimal control theory^45,46^. As a focus of future studies, we can introduce antigen proliferation into our current model, and consider its interaction with immune system, which could enable us to understand atopic dermatitis with immune memory formation.

Finally, our model would also enable us to address how immune responses change throughout our life as seen in the hygiene hypothesis. This hypothesis states that unhygienic experience (experience of lots of infection) during early childhood prevents allergic diseases; on the contrary, hygienic environments raise their risk^47^. The authenticity of this hypothesis is still controversial, but it suggests that discrimination of antigens can be influenced by all previous exposures to multiple antigens, that is, personal hygiene. Although we need to expand our model into the model that describes immune responses toward multiple antigens, our model might explain the difference in allergic risks based on individual antigen experiences.

## Methods

### T cell population dynamics with different properties of T cell activation

The model was extended to include the effect of dose-response of T cell activation on immune discrimination as

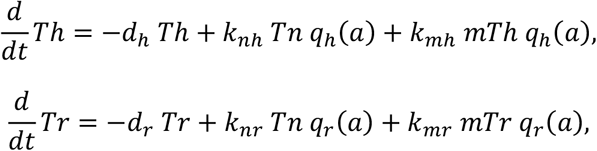

where *q_h_*(*a*) and *q_r_*(*a*) represent the dose-responses of Th and Tr cells, respectively, as described by the Hill equation:

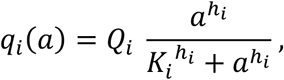

where *i* ∈ {*h*, *r*}, *Q_i_*, *K_i_* and *h_i_* indicate the amplitude, half-maximal effective antigen concentration, and Hill coefficient, respectively. Here, we considered three types of dose-response curves, namely linear, sigmoidal, and step-like types, where (*h_i_*, *Q_i_*, *K_i_*) = (1, 3500, 3000), (4, 10, 30), and (8, 10, 10), respectively. The same types of dose-response was applied for both Th and Tr cells in Fig. 5 (*q_h_*(*a*) = *q_r_*(*a*)), whereas different types of dose-response were considered in Fig. S1 and S2 (*q_h_*(*a*) = *q_r_*(*a*) or *q_h_*(*a*) ≠ *q_r_*(*a*)).

## Acknowledgements

We are grateful to Prof. Michiyuki Matsuda for providing the research environment. This study was supported in part by Moonshot R&D–MILLENNIA Program [grant number JPMJMS2024-9] by JST; Grant-in-Aid for Transformative Research Areas (B) [grant number 21H05170]; the Cooperative Study Program of Exploratory Research Center on Life and Living Systems (ExCELLS) [program number 19-102 to H.N.]. It was also supported by JSPS KAKENHI [grant number JP21J23680 to K.Y.] and a Grant-in-Aid for Young Scientists (B) [grant numbers 16K16147 and 19H04776 to H.N.] from the Japan Society for the Promotion of Science (JSPS).

## Author contribution

K.Y. and H.N. conceived the project and developed the method, K.Y. implemented the model simulation. K.Y. and H.N. wrote the manuscript.

## Data availability

No datasets were generated and analyzed during the current study.

## Supplementary Information

**Supplementary Table 1:** List of parameter values for simulation.

| T cell population dynamics |  |  |  |  |  |
| --- | --- | --- | --- | --- | --- |
| $d_h$ | 100 | $d_r$ | 10 | | |
| $k_{nh}$ | 1 | $k_{nr}$ | 0.01 | | |
| $k_{mh}$ | 4 | $k_{mr}$ | 0.4 | | |
| $d_{mh}$ | 0 | $d_{mr}$ | 0 | | |
| $Tn$ | 1 | $r_0$ | 0.01 | | |
| $r_s$ | 0.01 | $r_a$ | 0.01 | | |
| Predictive coding |  |  |  |  |  |
| $e_h$ | 0.0001 | $e_r$ | 0.0003 | | |
| $m_h$ | 1 | $m_r$ | 1 | | |
| $A_{max}$ | 100 | $K$ | 1 | | |
| Different dose-responses of T cell activation |  |  |  |  |  |
| Linear |  | Sigmoidal |  | Step-like |  |
| $h_i$ | 1 | $h_i$ | 4 | $h_i$ | 8 |
| $Q_i$ | 3500 | $Q_i$ | 10 | $Q_i$ | 10 |
| $K_i$ | 3000 | $K_i$ | 30 | $K_i$ | 10 |

**Supplementary Figure 1:**
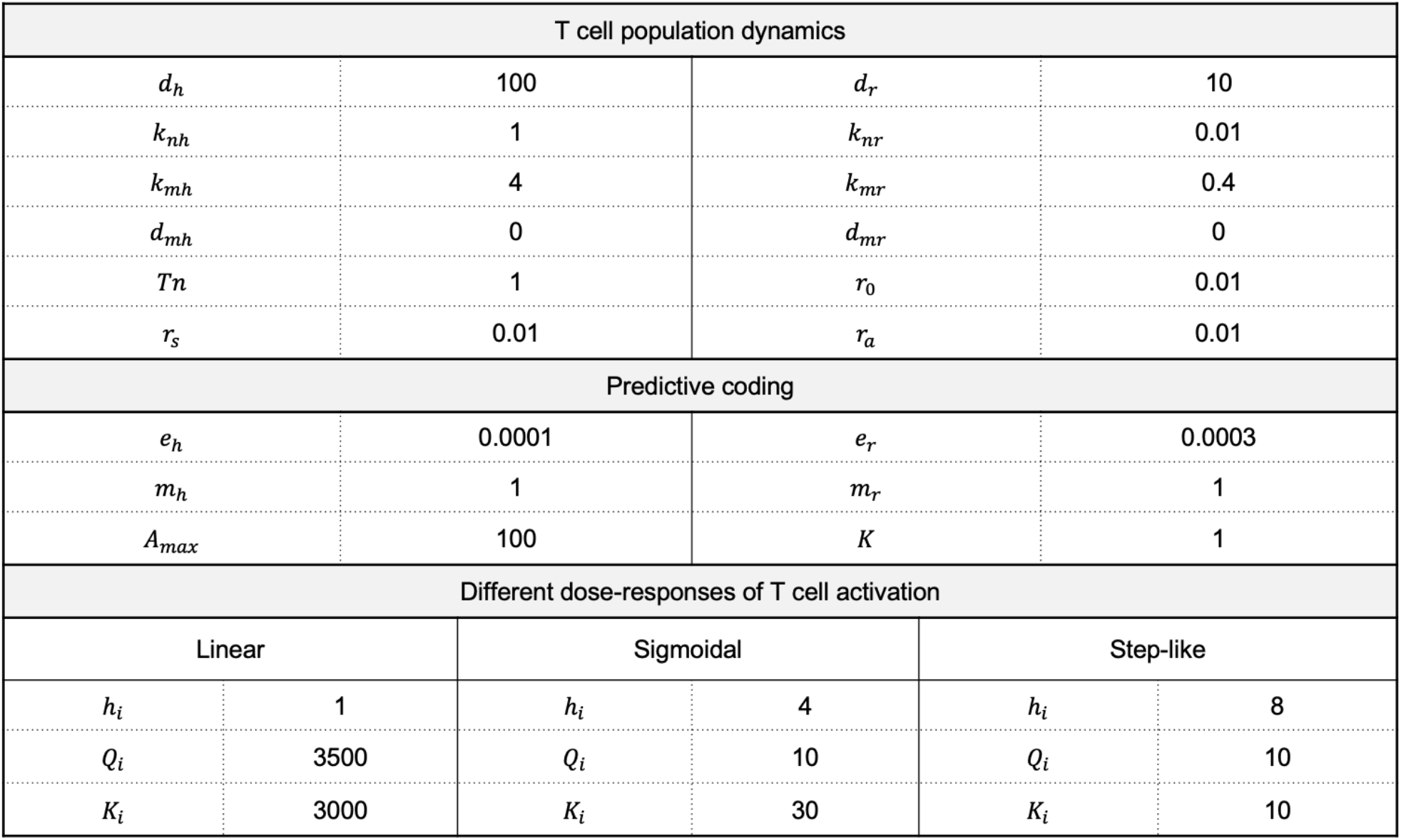
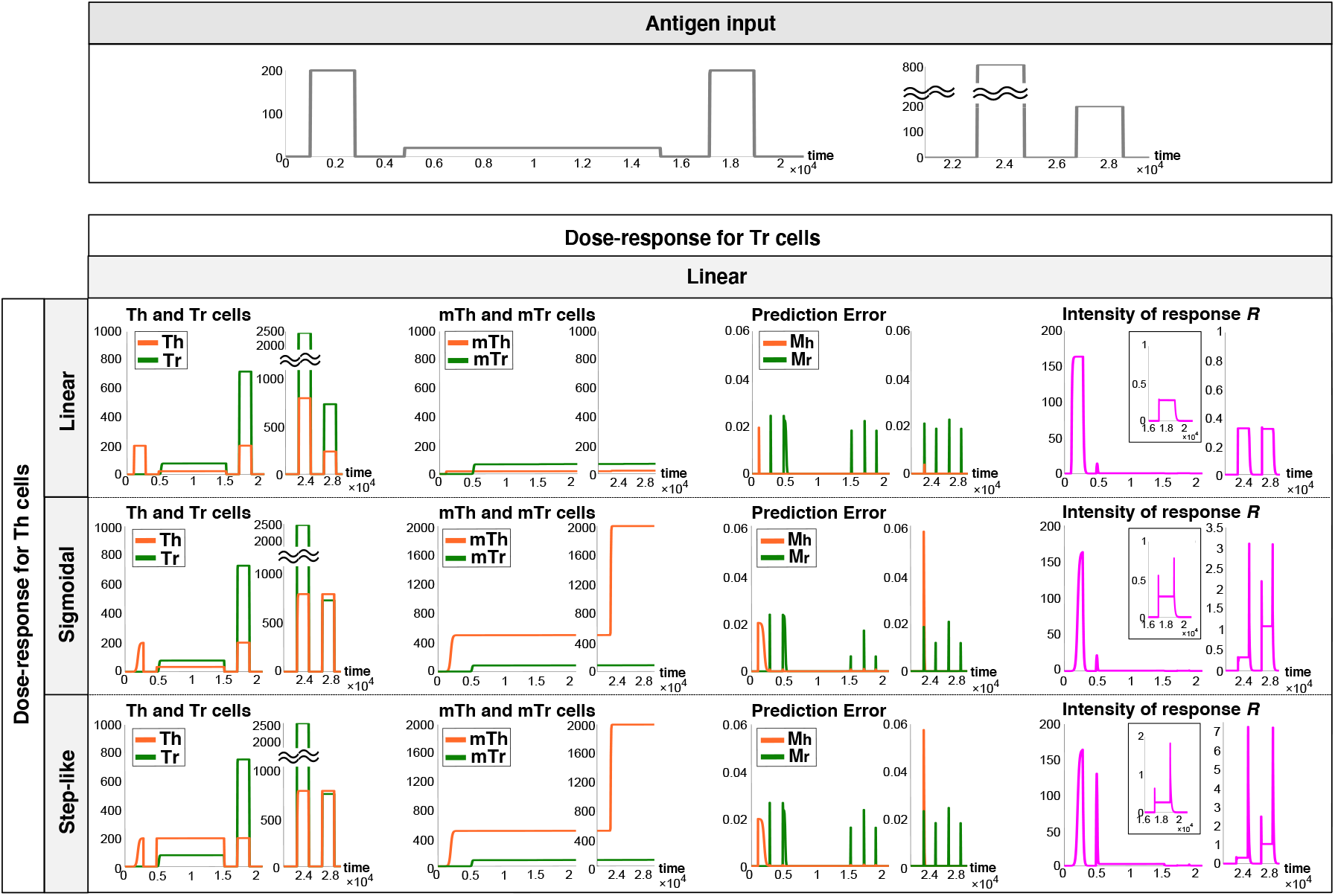
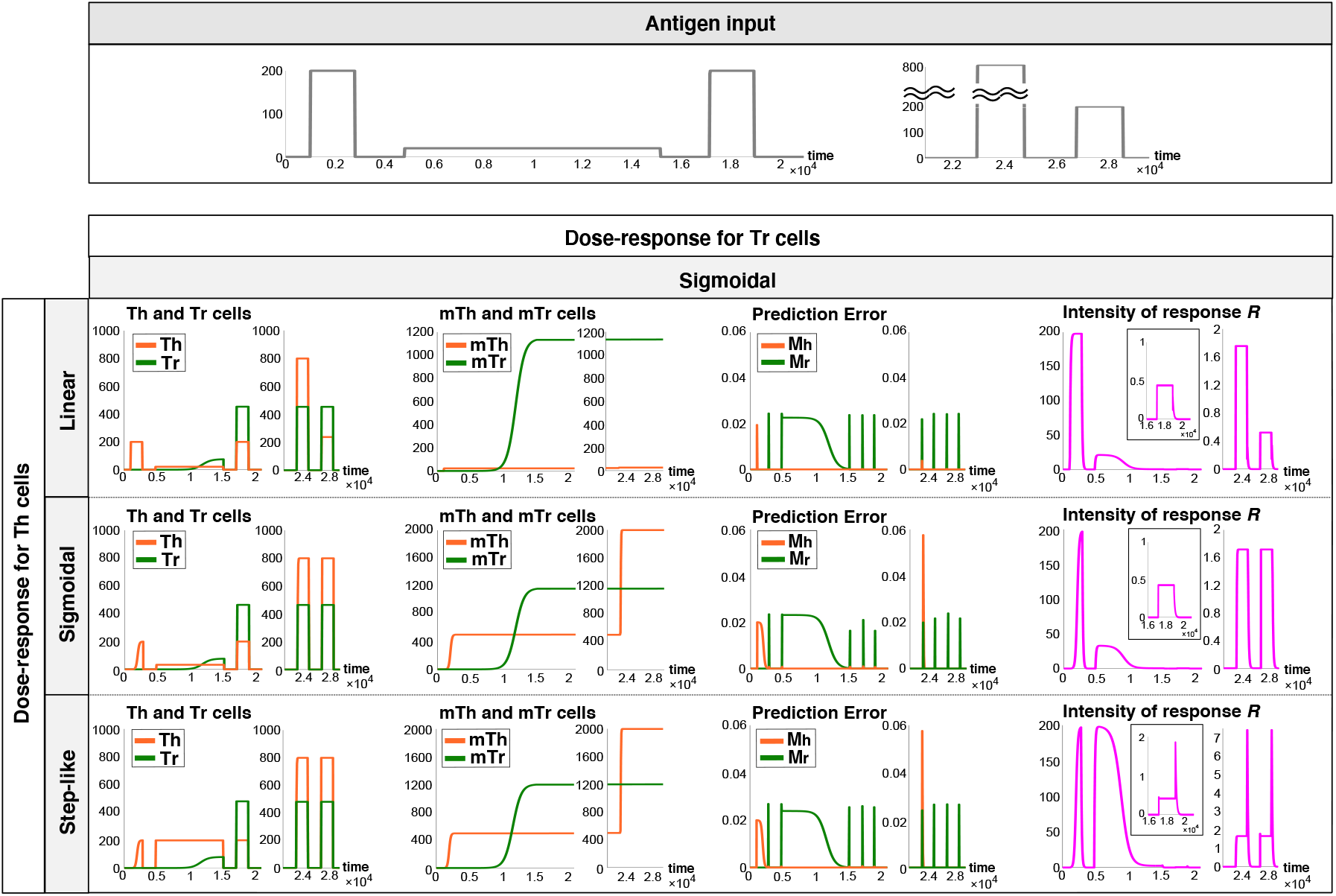
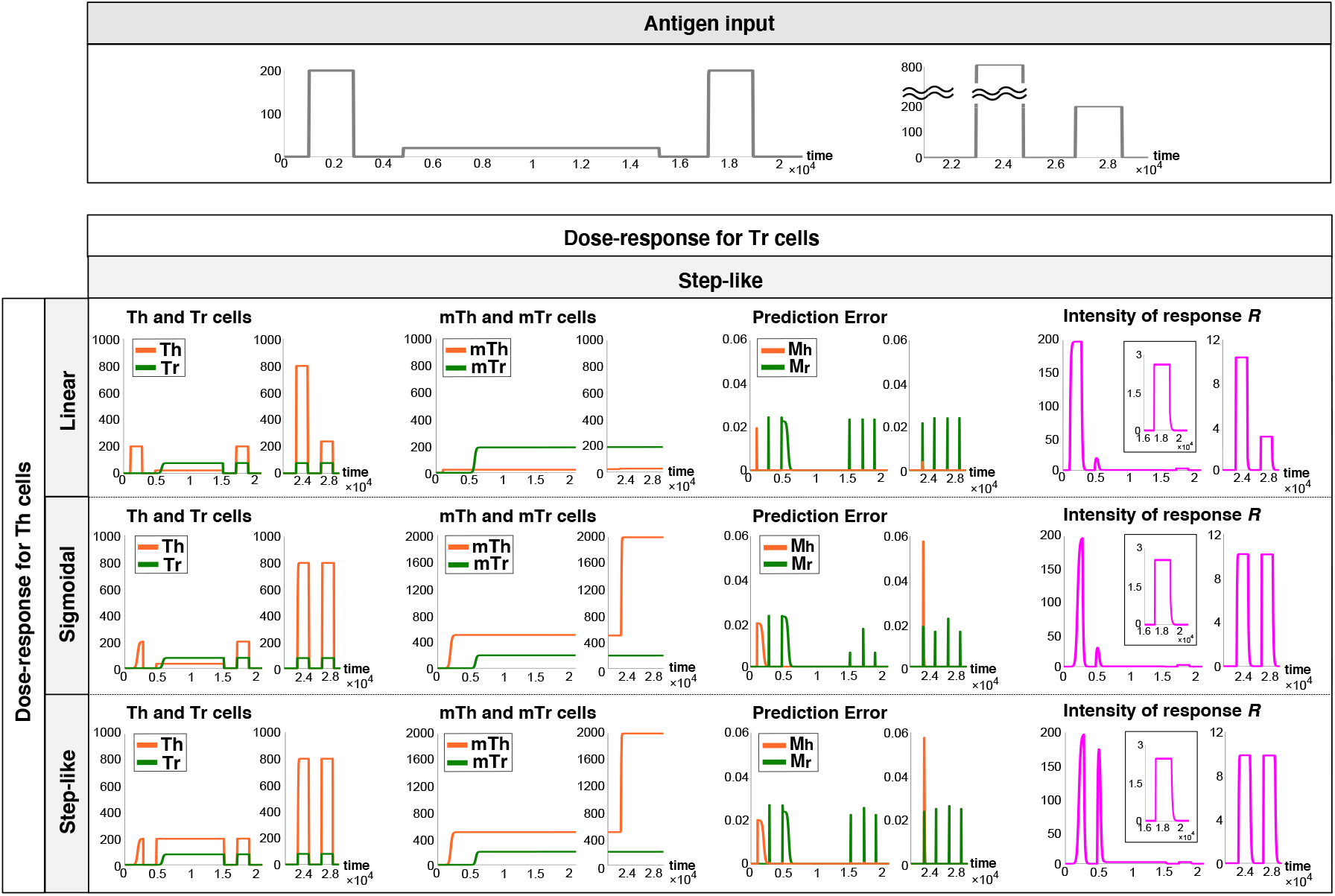
Simulations of therapeutic effect and its persistence under 9 combinations of dose-responses of T cell activation. Temporal changes in immune responses to a series of different antigen inputs (shown in each top panel) under all combinations of dose-response types for Th and Tr cell activation. The first antigen input was high enough for the induction of allergy, the second was applied for immunotherapy, and the third was for checking the therapeutic effect. The fourth antigen input was higher than the first one, and the fifth was applied to examine the effect of the fourth antigen input.

**Supplementary Figure 2:**
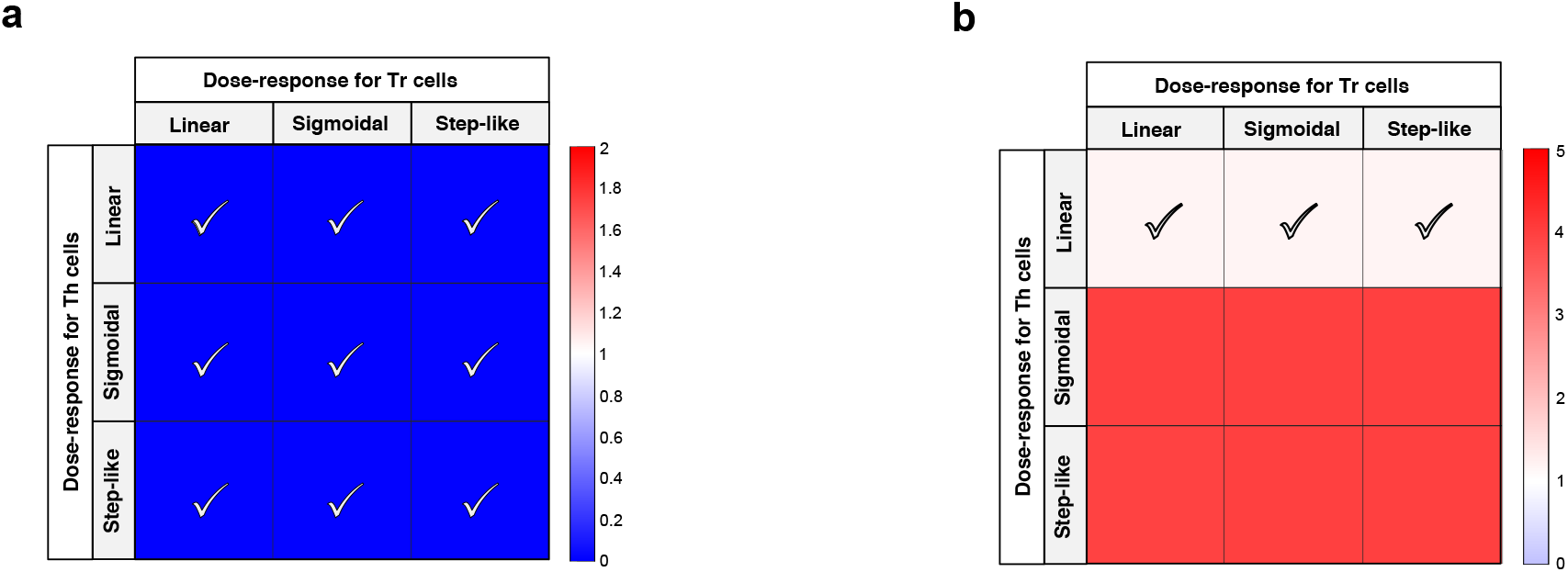
Therapeutic effect and its persistence in 9 combinations of doseresponses of T cell activation. **(a)** Heatmap showing therapeutic effect in all combinations of dose-response types of Th and Tr cell activation. The therapeutic effect was evaluated by ratio of the maximum intensity of response to the third antigen input to that to the first antigen input. The check marks indicate the cases in which the therapeutic effect was less than 1, that is, immunotherapy was successful. **(b)** Heatmap showing the persistence of immunotherapy in all combinations of dose-response types of Th and Tr cell activation. Persistence was quantified by ratio of the maximum intensity of response to the fifth antigen input to that to the third antigen input. Check marks indicate the cases where the quantified persistence was close to 1, that is, the effect of immunotherapy persisted after the fourth antigen input. In the calculation of the maximum intensity on each antigen input, we ignored the transient peaks of intensity at the initiation and end of antigen input on the numerical simulation.

## References

1. Evavold, B. D. & Allen, P. M. Separation of IL-4 production from Th cell proliferation by an altered T cell receptor ligand. Science (80-.). 252, (1991).

2. Dustin, M. L. T-cell activation through immunological synapses and kinapses. Immunological Reviews vol. 221 (2008).

3. Monks, C. R. F., Freiberg, B. A., Kupfer, H., Sciaky, N. & Kupfer, A. Three-dimensional segregation of supramolecular activation clusters in T cells. Nature 395, (1998).

4. Kappler, J. W., Roehm, N. & Marrack, P. T cell tolerance by clonal elimination in the thymus. Cell 49, (1987).

5. Kisielow, P., Blüthmann, H., Staerz, U. D., Steinmetz, M. & Von Boehmer, H. Tolerance in T-cell-receptor transgenic mice involves deletion of nonmature CD4+8+ thymocytes. Nature 333, (1988).

6. Anderson, M. S. et al. Projection of an immunological self shadow within the thymus by the aire protein. Science (80-.). 298, (2002).

7. Luckheeram, R. V., Zhou, R., Verma, A. D. & Xia, B. CD4 +T cells: Differentiation and functions. Clinical and Developmental Immunology vol. 2012 (2012).

8. Crotty, S. Follicular Helper CD4 T cells (T FH). Annu. Rev. Immunol. 29, (2011).

9. Zhu, J. & Paul, W. E. CD4 T cells: Fates, functions, and faults. Blood 112, (2008).

10. Vignali, D. A. A., Collison, L. W. & Workman, C. J. How regulatory T cells work. Nature Reviews Immunology vol. 8 (2008).

11. Sakaguchi, S., Yamaguchi, T., Nomura, T. & Ono, M. Regulatory T Cells and Immune Tolerance. Cell vol. 133 (2008).

12. Chowdhury, D. & Lieberman, J. Death by a thousand cuts: Granzyme pathways of programmed cell death. Annual Review of Immunology vol. 26 (2008).

13. Brincks, E. L. et al. Antigen-Specific Memory Regulatory CD4 + Foxp3 + T Cells Control Memory Responses to Influenza Virus Infection. J. Immunol. 190, (2013).

14. Gasper, D. J., Tejera, M. M. & Suresh, M. CD4 T-cell memory generation and maintenance. Crit. Rev. Immunol. 34, (2014).

15. Kaech, S. M., Wherry, E. J. & Ahmed, R. Effector and memory T-cell differentiation: Implications for vaccine development. Nature Reviews Immunology vol. 2 (2002).

16. Harrington, L. E., Janowski, K. M., Oliver, J. R., Zajac, A. J. & Weaver, C. T. Memory CD4 T cells emerge from effector T-cell progenitors. Nature 452, (2008).

17. Garcia, S., DiSanto, J. & Stockinger, B. Following the development of a CD4 T cell response in vivo: From activation to memory formation. Immunity 11, (1999).

18. Canonica, G. W. et al. Sublingual immunotherapy: World Allergy Organization position paper 2013 update. World Allergy Organization Journal vol. 7 (2014).

19. Noon, L. PROPHYLACTIC INOCULATION AGAINST HAY FEVER. Lancet 177, (1911).

20. Pfaar, O. et al. Guideline on allergen-specific immunotherapy in IgE-mediated allergic diseases. Allergo J. Int. 23, (2014).

21. Böhm, L. et al. IL-10 and Regulatory T Cells Cooperate in Allergen-Specific Immunotherapy To Ameliorate Allergic Asthma. J. Immunol. 194, (2015).

22. Shamji, M. H. & Durham, S. R. Mechanisms of immunotherapy to aeroallergens. Clinical and Experimental Allergy vol. 41 (2011).

23. Radulovic, S., Jacobson, M. R., Durham, S. R. & Nouri-Aria, K. T. Grass pollen immunotherapy induces Foxp3-expressing CD4+CD25+ cells in the nasal mucosa. J. Allergy Clin. Immunol. 121, (2008).

24. Rao, R. P. N. & Ballard, D. H. Predictive coding in the visual cortex: A functional interpretation of some extra-classical receptive-field effects. Nat. Neurosci. 2, (1999).

25. Friston, K. The free-energy principle: A unified brain theory? Nature Reviews Neuroscience vol. 11 (2010).

26. Friston, K. & Kiebel, S. Predictive coding under the free-energy principle. Philos. Trans. R. Soc. B Biol. Sci. 364, (2009).

27. Friston, K. J., Daunizeau, J., Kilner, J. & Kiebel, S. J. Action and behavior: A free-energy formulation. Biol. Cybern. 102, (2010).

28. Turner, M. D., Nedjai, B., Hurst, T. & Pennington, D. J. Cytokines and chemokines: At the crossroads of cell signalling and inflammatory disease. Biochimica et Biophysica Acta - Molecular Cell Research vol. 1843 (2014).

29. Sturm, G. J. et al. EAACI guidelines on allergen immunotherapy: Hymenoptera venom allergy. Allergy Eur. J. Allergy Clin. Immunol. 73, (2018).

30. Cox, L. et al. Allergen immunotherapy: A practice parameter third update. J. Allergy Clin. Immunol. 127, (2011).

31. Barni, S. et al. Immunoglobulin E (IgE)-mediated food allergy in children: Epidemiology, pathogenesis, diagnosis, prevention, and management. Medicina (Lithuania) vol. 56 (2020).

32. Aleksic, M. et al. Dependence of T Cell Antigen Recognition on T Cell Receptor-Peptide MHC Confinement Time. Immunity 32, (2010).

33. Čemerski, S. et al. The Stimulatory Potency of T Cell Antigens Is Influenced by the Formation of the Immunological Synapse. Immunity 26, (2007).

34. Evavold, B. D., Sloan-Lancaster, J. & Allen, P. M. Tickling the TCR: selective T-cell functions stimulated by altered peptide ligands. Immunology Today vol. 14 (1993).

35. Turner, P. J. et al. Fatal Anaphylaxis: Mortality Rate and Risk Factors. J. Allergy Clin. Immunol. Pract. 5, (2017).

36. Ratajczak, W., Niedźwiedzka-Rystwej, P., Tokarz-Deptuła, B. & DeptuŁa, W. Immunological memory cells. Central European Journal of Immunology vol. 43 (2018).

37. Rosenblum, M. D., Way, S. S. & Abbas, A. K. Regulatory T cell memory. Nature Reviews Immunology vol. 16 (2016).

38. Bianchi, D. W., Zickwolf, G. K., Weil, G. J., Sylvester, S. & Demaria, M. A. Male fetal progenitor cells persist in maternal blood for as long as 27 years postpartum. Proc. Natl. Acad. Sci. U. S. A. 93, (1996).

39. Nelson, J. L. The otherness of self: Microchimerism in health and disease. Trends in Immunology vol. 33 (2012).

40. Sanchez, A. M., Zhu, J., Huang, X. & Yang, Y. The Development and Function of Memory Regulatory T Cells after Acute Viral Infections. J. Immunol. 189, (2012).

41. Hara, A. & Iwasa, Y. When is allergen immunotherapy effective? J. Theor. Biol. 425, (2017).

42. Groß, F., Metzner, G. & Behn, U. Mathematical modeling of allergy and specific immunotherapy: Th1-Th2-Treg interactions. J. Theor. Biol. 269, (2011).

43. Sontag, E. D. A Dynamic Model of Immune Responses to Antigen Presentation Predicts Different Regions of Tumor or Pathogen Elimination. Cell Syst. 4, (2017).

44. Pradeu, T., Jaeger, S. & Vivier, E. The speed of change: Towards a discontinuity theory of immunity? Nature Reviews Immunology vol. 13 (2013).

45. Domínguez-Hüttinger, E. et al. Mathematical modeling of atopic dermatitis reveals “double-switch” mechanisms underlying 4 common disease phenotypes. J. Allergy Clin. Immunol. 139, (2017).

46. Christodoulides, P. et al. Computational design of treatment strategies for proactive therapy on atopic dermatitis using optimal control theory. Philos. Trans. R. Soc. A Math. Phys. Eng. Sci. 375, (2017).

47. Yazdanbakhsh, M., Kremsner, P. G. & Van Ree, R. Immunology: Allergy, parasites, and the hygiene hypothesis. Science vol. 296 (2002).

